# The RNA content of extracellular vesicles from *PRPF31*^+/−^ hiPSC-RPE show potential as biomarkers of retinal degeneration

**DOI:** 10.1101/2022.04.05.487197

**Authors:** Heran Getachew, Sudeep Mehrotra, Tarandeep Kaur, Rosario Fernandez-Godino, Eric A. Pierce, Marcela Garita-Hernandez

## Abstract

Retinitis pigmentosa (RP), is the most common inherited retinal degeneration (IRD), leading to vision loss via dysfunction and death of photoreceptor cells and retinal pigment epithelium (RPE). Mutations in the pre-mRNA processing factor 31 (*PRPF31*) gene are associated with autosomal dominant RP, impairing RPE function. While adeno-associated virus (AAV)-mediated gene therapy shows promise for treating IRDs, the slow progression of these diseases often makes timely measurement of clinical efficacy challenging. Extracellular vesicles (EVs) are lipid enclosed vesicles secreted by cells and their RNA contents are being explored as circulating biomarkers of cancer and other diseases. We hypothesize that EV RNAs could serve as biomarkers of the health status of the neural retina and RPE health. To test this, we used *PRPF31*^+/+^ and *PRPF31*^+/−^ human-induced pluripotent stem cell (hiPSC)-derived RPE (hi-RPE) to investigate the RNAs contained in RPE-derived EVs, and how they change in disease. We also compared the RNA contents of RPE-EVs with the RNAs contained in the hi-RPE cells themselves. We found that EVs from mutant *PRPF31*^+/−^ hi-RPE cells have distinct RNA profiles compared to those from control cells, suggesting EV RNA contents change during disease and could serve as biomarkers for retinal degeneration.

## Introduction

Inherited retinal degenerations (IRDs) are significant contributors to vision loss in both children and adults ^1,2^. Among these, rod-cone degeneration, commonly known as retinitis pigmentosa (RP), is the most prevalent form of IRD, affecting roughly 1 in 2,500 individuals globally ^3^. The pathology of IRD and RP includes degeneration of the retinal pigment epithelium (RPE) and/or neural retina, which lead to vision loss ^4^. Mutations in pre-RNA processing factors (PRPFs) are among the primary causes of autosomal dominant RP ^5,6^, with mutations in *PRPF31* being the most frequent, accounting for approximately 10% of cases ^7^. Notably, mutations in *PRPF31* lead to dominant disease through haploinsufficiency ^8–12^.

Research using mouse models of *PRPF31*-associated RP have shown that haploinsufficiency of *PRPF31* causes cell autonomous defects in RPE function, including reduced phagocytosis of photoreceptor outer segments, which is crucial for RPE and retinal health ^13–17^. Similarly, *PRPF31^+/−^* mutant human induced pluripotent stem cell (hiPSC) derived RPE (hiPSC-RPE) cells exhibit reduced phagocytosis function, shorter cilia, and reduced barrier function of the RPE ^16–18^. Gene augmentation therapies have shown promise in addressing the consequences of haploinsufficiency of *PRPF31* gene ^17,19^. Adeno-associated virus (AAV) mediated gene augmentation therapy was able to restore normal function to *PRPF31^+/−^* hiPSC-RPE cells, suggesting its potential use to treat *PRPF31*-associated RP ^17,18^. Gene augmentation has also been shown to prevent retinal degeneration in a somatic mouse model of *PRPF31* deficiency ^20^.

Genetic therapies including AAV – mediated gene therapy have shown great promise for the treatment of additional genetic forms of IRD ^21–25^. Given the slow progression in many IRDs, central vision can be retained until late stages of disease making challenging to demonstrate the benefit of treatment during clinical trials. It can take years for improvement in clinical outcome measures such as visual acuity and visual fields to become detectable ^26^. Therefore, surrogate markers of retinal health could greatly enhance the monitoring of IRD patients during trials. One promising type of biomarker is RNAs contained within extracellular vesicles (EVs). EVs are cell-derived membranous structures (40–1,000 nm in diameter) ^27,28^ that carry a diverse cargo including lipids, proteins, RNA and DNA ^29,30^. Present both locally and systemically, EVs are believed to facilitate intercellular communication by transferring their protein and RNA between cells ^31–33^. EVs are secreted from nearly all cell types and are found in various biofluids, including plasma, urine, and the vitreous and aqueous humors of the eye ^34^. Their potential as biomarkers is supported by their ability to mirror the health status of their source cells ^29,35–39^.

Liquid biopsies of EV contents are proving effective in monitoring tumor burden and minimal residual disease across various cancers ^40^. Early research suggests that the contents of retina and RPE-derived EVs may reflect retinal health status ^41,42^, although systematic studies of RNA contents in these EVs are still lacking. Several studies have explored the potential of RNA in EVs from the vitreous, aqueous humor, and serum as biomarkers for conditions such as uveal melanoma, age-related macular degeneration, and glaucoma ^43–47^. Additionally, the protein content of EVs produced by the retinal pigment epithelium (RPE) has also been investigated ^41,48^. Research involving RPE cells derived from swine primary cultures and retinal organoids has demonstrated their ability to release EVs ^48^. Notably, proteomic analyses of these EVs have linked their content to age-related macular degeneration ^49^. However, their precise function in retinal pathology remains largely unexplored.

In order to test the hypothesis that EV RNAs can be used as biomarkers for *PRPF31*-associated RP as well as other IRDs affecting the RPE, it is crucial to define the RNA profiles of RPE-derived EVs. In this study, we characterized the RNA profiles of EVs from control *PRPF31*^+/+^ and mutant *PRPF31*^+/−^ hiPSC-RPE cells. We also compared these profiles with the RNA transcriptomes of the originating RPE cells. Our findings reveal that the RNA cargo in EVs mirrors that of the originating RPE cells, with distinct differences reflecting disease perturbation and regulatory changes in genes associated with RPE de-differentiation and function. These results support the potential of RNA biomarkers in assessing RPE health status.

## Materials and Methods

### Generation of hiPSC-derived RPE cell cultures

Wildtype *PRPF31* and mutant *PRPF31^+/−^* hiPSC were previously engineered using CRISPR Cas9 genome editing ^17,50^. These hiPSC were maintained on growth-factor-reduced Matrigel (BD Biosciences) coated plates and their pluripotency state was confirmed by expression of pluripotency markers OCT4, NANOG and SSEA4 (**Supplementary Figure 1**). The differentiation capacity of the lines was assessed with the spontaneous differentiation Embryoid Body (EB) model, which demonstrated *PRPF31^+/−^*and its isogenic control hiPSC line were capable of differentiating into the three germ layers: Ectoderm, mesoderm and endoderm (**Supplementary Figure 1**). For the directed differentiation towards RPE, *PRPF31^+/+^* and mutant *PRPF31^+/−^* hiPSC were left in culture until an approximate 90% confluence was reached (Day 0) and differentiated as previously described ^17^. Briefly, hiPSC were treated with a combination of Noggin, Dickkopf-1 (Dkk-1), Insulin Growth Factor 1, Nicotinamide, Activin A, basic Fibroblast Growth Factor, SU-5402, and CHIR99021 to induce the directed differentiation of hiPSCs into RPE cells. On day 30, cells were enzymatically digested with TrypLE^TM^ Express (12604013, Thermo Fisher, Waltham, MA, USA), strained through a 40-mm filter, and seeded at a density of 1X10^5^ cells/cm^2^ onto Matrigel-coated 6-well plates in X-VIVO 10 media. After 60 days in culture, hi-RPE cells were seeded onto non-growth factor reduced Matrigel-coated 24-mm Transwell® with 0.4-μm pore polyester membrane inserts (3450, Corning, Bedford, MA, USA) at a density of 700,500 cells per well. The cells were matured for 4 weeks on Transwells® before conditioned media was collected (**Figure 1**). Apical and basal conditioned media was collected every other day for 12 weeks and stored at -80°C.

**Figure 1.**
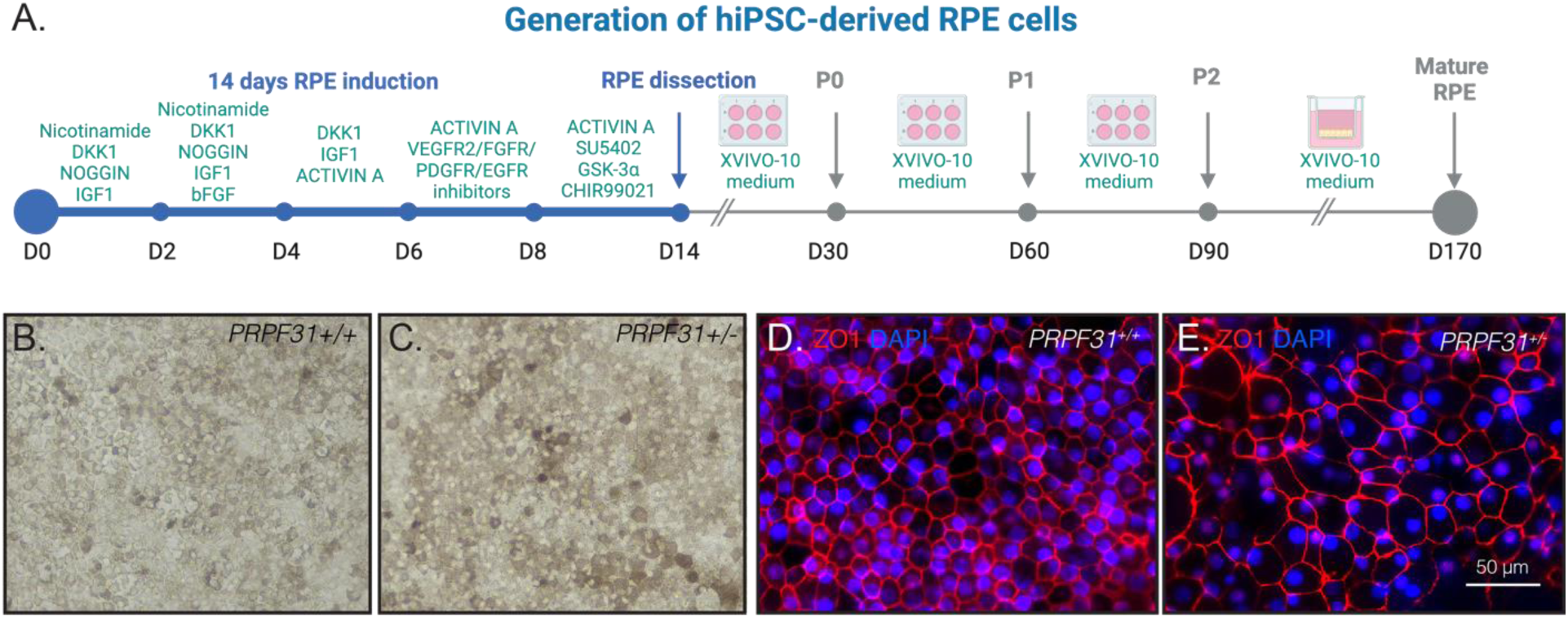
Generation of hiPSC-derived RPE cells. (A) Schematic process of the directed differentiation of PRPF31^+/+^ and PRPF31^+/−^ hiPSC into RPE cells (B–C) Phase contrast images of PRPF31^+/+^ and PRPF31^+/-^ hiPSC-RPE showing the cobblestone morphology of the pigmented RPE cells. (D-E) Confocal images of PRPF31^+/+^ and PRPF31^+/-^ hiPSC-RPE cells immunoreactive for the tight junction marker ZO1.

### EV isolation

The conditioned media from serum-free RPE cell cultures was thawed on ice and centrifuged 3000 × g for 15 minutes to remove cells and cell debris. EVs were isolated using ExoQuick-TC (EXOTC50A-1, System Biosciences, Palo Alto, CA, USA) according to manufacturer instructions. Briefly, the conditioned was mixed with ExoQuick-TC and incubated at 4°C for at least 12 hours. Then, ExoQuick-TC/conditioned media mixture was centrifuged at 1500 × g for 30 minutes and the supernatant was discarded. A final spin was performed at 1500 × g for 5 minutes to remove any residual fluid. The EV pellet was resuspended in 100 μL of 1X particle free phosphate-buffered saline (PBS) and stored at -80°C (**Figure 1A**). The isolated EVs, ranging in size from 40 to 1000 nm, were characterized as EVs using EM imaging. Since this study concentrates on the RNA content as potential biomarkers of the isolated EVs, and because non-vesicular extracellular particles lack RNA, no further separation was deemed necessary in accordance with the latest Minimal Information for Studies of Extracellular Vesicles (MISEV) recommendations ^51^.

### Nanoparticle tracking analysis (NTA)

The size distribution and concentration of EVs was determined using a NanoSight NS300 instrument (Malvern Instruments Ltd, Malvern, UK) as previously described (Mass SL 2015). The EV suspensions were diluted 1:100 in PBS, and 1 mL was used for NTA. Size and diffusion of nanoparticles was tracked, recorded, and analyzed using NTA software 3.2. Detection threshold was set for each sample individually in a way to meet manufacturer recommended quality standards.

Transmission Electron Microscopy TEM)

EV morphologies were visualized with transmission electron microscopy (TEM) following a procedure adapted from Théry et al ^52^. Briefly, the EVs were lyophilized and fixed for 1 hour at room temperature in 2% paraformaldehyde in 0.1 mol/L sodium phosphate buffer (Electron Microscopy Sciences, Hatfield, Pa). Five microliters of the exosome were absorbed onto Formvar/carbon-coated electron microscopy grids (Electron Microscopy Sciences) for 20 minutes. The grids were rinsed 8 times in 0.1 mol/L sodium phosphate buffer and then incubated in 1% glutaraldehyde in 0.1 mol/L sodium phosphate buffer (Electron Microscopy Services) for 5 minutes. After rinsing 8 times in deionized water, grids were contrasted in uranyl-oxalate solution (pH 7; Electron Microscopy Services) for 5 minutes and transferred to uranyl acetate-methyl cellulose solution for 5 minutes. The grids were blotted on filter paper and air-dried before imaging. The EVs were observed with an FEI Tecnai G2 Spirit transmission electron microscope (FEI, Hillsboro, Ore) at an accelerating voltage of 100 kV interfaced with an AMT XR41 digital CCD camera (Advanced Microscopy Techniques, Woburn, Mass) for digital TIFF file image acquisition.

### RNA extraction and purification

Total RNA was isolated from 200µl of EV using Seramir Exosome RNA amplification kit following manufacturer’s instructions. Purified RNA was quantified using Agilent RNA 6000 Pico kit in a Bioanalyzer.

### miRNA Sequencing

RNA was isolated from the EVs using the ExoQuick RNA column purification kit (EQ808A-1, System Biosciences, Palo Alto, CA, USA). Purified RNA was quantified using a Bioanalyzer (Agilent 2100 Bioanalyzer) and the Agilent RNA 6000 Pico kit (5067-1513, Agilent, Santa Clara, CA, USA). miRNA libraries were prepared from the isolated EV RNA and a water control using the TruSeq Small RNA Library Preparation Kit (RS-200-0012, Illumina, San Diego, CA, USA) using the manufacturer instructions with the modification of performing the adaptor ligation in the presence of 15% polyethylene glycol in order to reduce bias by driving the ligation reactions towards completion as determined by the Extracellular RNA Communication Consortium^53^. Adapter dimers were removed by gel purification and the purified cDNA libraries were pooled and normalized 2nM. Nuclease free water was added to the samples as a control during sequencing. The pooled libraries were then sequenced single-read, 50bp (1 × 50) on an Illumina MiSeq using version 3 reagents at the Ocular Genomic Institute, MEE, Boston, MA (USA).

### miRNA Sequencing Analysis

Base quality per sequence cycle was checked with FastQC. An average minimum mean base quality above Q30 (Phred scale) was observed using MultiQC across the read length for all samples. The sample replicates were sequenced across four different sequencing runs. Each sequenced run was independently quality checked. The files were merged and quality checked again for any biases. Illumina TruSeq small RNA adapters were scanned on the 5’ and 3’ ends and trimmed using Cutadapt(v.3.4). Post trimming, minimum length of 17 bp was maintained. On average 1.5 million reads were observed across all the samples. Samples less then 1 million reads were dropped. Following normalization using edgeR (v3.30), multidimensional scaling (MDS) plot with biological coefficient of variations (BCV) were further used to identify outlier samples and were removed from all downstream analysis. The standard miRDeep2 (v2.0.1.2) was used for the analysis. Briefly, mapper module (mapper.pl) was used for read alignment against the reference human genome (GRCh38). Identification and quantification of each miRNA was performed by the quantifier module (quantifier.pl). Precursor and mature miRNAs sequences of human miRBase (miRNA repository) was used as reference sequence. For novel miRNA prediction, mature miRNA of closely related species (*Gorilla gorilla, Pongo pygmaeus, Pan troglodytes and Pan paniscus*) were pooled together for alignment and novel miRNA discoveries. The counts of miRNA identified and reported in the water sample were used to penalize the identified miRNA in other sample replicates. Samples with alignment rate of <1% were further removed. The Trimmed Mean of the M-values (TMM) from the edgeR package was used for normalized within and across sample replicates. Post normalization, counts per million (CPM) values were to filter and identify “expressed” miRNA across samples replicates. The standard settings in the edgeR package used for differential analysis. Gene set enrichment analysis; Gene Ontology (GO) pathways (KEGG) and diseases was performed using miEAA a web-based application which performs rich functionality in terms of miRNA categories and determines if a category is significantly enriched in a given miRNA set with respect to a reference^54,55^.

### EV Poly-A RNA Sequencing

cDNA was prepared from EV RNA using the Single Cell RNA kit SMART-*Seq V4* Ultra Low Input RNA Kit (634890, Takara, Mountain View, CA, USA) according to manufacturer instructions using 1ng input. Next, poly-A libraries were prepared using the SMARTer ThruPLEX DNA-seq Kit (R400675, Takara, Mountain View, CA, USA). Each library was normalized to 1.5 nM, pooled and 1% PhiX was added. Libraries were sequenced for 101 cycles (2 × 101bp (PE)) using the NovaSeq 6000 (Illumina) at the Ocular Genomic Institute, MEE, Boston, MA (USA).

### RPE poly-A RNA Sequencing

RNA was extracted from hiPSC-RPE cells and Spike-in RNA standards, sequins, were used as a reference in all samples ^56^. Different sets of sequin mixtures were used, one for wildtype *PRPF31^+/+^* and another set for mutant *PRPF31^+/−^*samples. Libraries were prepared using the NEBNext Ultra II Directional RNA Library Prep with Sample Purification Beads (E7765S, New England Biolabs, Ipswich, MA, USA), NEBNext Poly(A) mRNA Magnetic Isolation Module (E7490S, New England Biolabs, Ipswich, MA, USA), and NEBNext Multiplex Oligos for Illumina (E6440S, New England Biolabs, Ipswich, MA, USA). Libraries were normalized, pooled, and sequenced as described above.

### Poly-A RNAseq Analysis of RPE cells and its EVs

Read and sample level QC was performed. Read quality was assessed with FastQC (https://www.bioinformatics.babraham.ac.uk/projects/fastqc/) and MultiQC. An in-house Perl script was used to check and filter out reads with presence of adapter and ambiguous character, ‘N’. The Bowtie2 aligner was used to identify ribosomal contamination. Reads aligned to rRNA reference sequences were dropped from all downstream analysis using SAMtools, resulting in high quality (HQ) reads. The STAR aligner was used to align PE reads to a mouse reference in two-pass mode within the sample and across replicates for each sample set. Feature Counts from the Subread package (v2.0.3), was used to generate gene expression matrix with the following non-default settings: reads must be paired, both the pairs must be mapped, use only uniquely mapped reads, multi-mapped reads are not counted, chimeric reads were not counted, and strand specificity turned on. Anaquin was used to further evaluate alignment sensitivity and gene expression ^57^. Here sensitivity indicates the fraction of annotated regions covered by alignment. The Picard tools (http://broadinstitute.github.io/picard/) and RSeQC were used to calculate mean fragment length. The approach implemented in Kallisto was used to covert raw reads to TPM values. An average TPM of the lowest Sequins between test and control samples was calculated and used as cutoff. DESeq2 was used for differential gene expression analysis. RNA-seq analysis is described in detail in Mehrotra et al. ^58^. g:Profiler (version e101_eg48_p14_baf17f0) with a custom background gene set was used for gene set enrichment analysis (GSEA), pathway enrichment and transcription factor binding site analysis.

### Reverse transcription and quantitative real-time polymerase chain reaction

hi-RPE monolayers scraped off the plate into a 1.5-ml tube. Cells were washed twice in ice cold PBS and centrifuged at 13.000 × *g* for 1 min to obtain a dry pellet and store them at -80C. At the time of the analysis, the pellet was resuspended in RLT plus buffer and RNA extraction was performed according to manufacturer’s protocol (RNeasy Micro Kit; Qiagen). Reverse transcription was performed using SuperScript VILO cDNA Synthesis Kit (Life Technologies, Thermo Fisher Scientific). Quantitative real-time PCRs (RT-qPCR), reactions were performed with TaqMan^®^ Fast Advanced Master Mix and validated TaqMan^®^ Gene Expression Assays (Life Technologies, Thermo Fisher Scientific) on QuantStudio™3 Real-Time instrument (Applied Biosystems) for 40 cycles. Samples were run in quadruplicates, and expression levels were normalized using 18S as housekeeping gene in the case of poly-A RNAs and hsa-miR26a-5p for the miRNAs. The differential expression of the poly-A RNA and miRNA were expressed as fold change (2^ (-ΔΔCt). The relative expression between PRPF31^+/−^ hi-RPE and PRPF31^+/+^ hi-RPE cells was compared by *Students t-test* and p value ≤ 0.05 was considered as significant. A complete list of the TaqMan Gene Expression and miRNA Assays used are provided in **Supplementary Table S9 and S10**.

## Results

### Derivation and characterization of human RPE from human induced pluripotent stem cells haploinsufficient for *PRPF31*

We previously generated hiPSCs harboring a 10 bp deletion in exon 7 of the *PRPF31* gene using CRISPR/Cas9-mediated genome editing ^17^. The resulting *PRPF31*^+/-^ mutant and isogenic wild-type control hiPSCs displayed typical ES cell-like colony morphology and expressed pluripotency markers OCT4, SSEA4, NANOG, and SOX2 (**Supplementary Figure 1**). Both the *PRPF31*^+/−^ mutant and isogenic control hiPSCs did not exhibit any chromosomal abnormalities and were able to differentiate into the three embryonic germ cell layers evaluated by the spontaneous *in vitro* differentiation EB model (**Supplementary Figure 1**). The *PRPF31*^+/-^ mutant and isogenic control hiPSC lines were differentiated into RPE using established methods (**Figure 1A**)^17,59^. At Passage (P) 2, RPE cells differentiated from *PRPF31*^+/−^ mutant and *PRPF31*^+/+^ control hiPSCs display the typical cobblestone morphology pigmented cells (**Figure 1B-C**). They organized in monolayer polarized cultures and abundantly expressed the Zonula occludens (ZO1) tight junction associated protein (**Figure 1D-E**). We have previously shown that these *PRPF31^+/−^*mutant RPE cells produce reduced levels of PRPF31, demonstrate defects in phagocytosis function, cilia formation and barrier function compared to RPE from the isogenic control hiPSCs ^16,17^. Further, the *PRPF31*^+/-^ mutant RPE cells appear to have irregular shapes as shown by ZO1 staining compared to *PRPF31*^+/+^ control RPE cells, as recently reported for RPE cells differentiated from patient-derived hiPSC lines harboring either a duplication in exon 8 (c.709_734dup) or a deletion in exon 4 (c.269_273del) in the *PRPF31* gene^18^.

### *PRPF31* haploinsufficiency does not impact the number or physical properties of EVs produced by the source hiPSC-RPE cells

To investigate the effects of *PRPF31* haploinsufficiency and RPE dysfunction on the EVs produced by RPE cells, 12 wells each of differentiated *PRPF31*^+/+^ and *PRPF31*^+/−^ iPSC-RPE cells were matured on Transwell® inserts for 4 weeks. Once the hiPSC-RPE were mature, the apical and basal conditioned media was collected for 12 weeks. EVs were isolated from the conditioned media (CM) of the RPE monolayers, as described (**Figure 2A**) ^60^.

**Figure 2.**
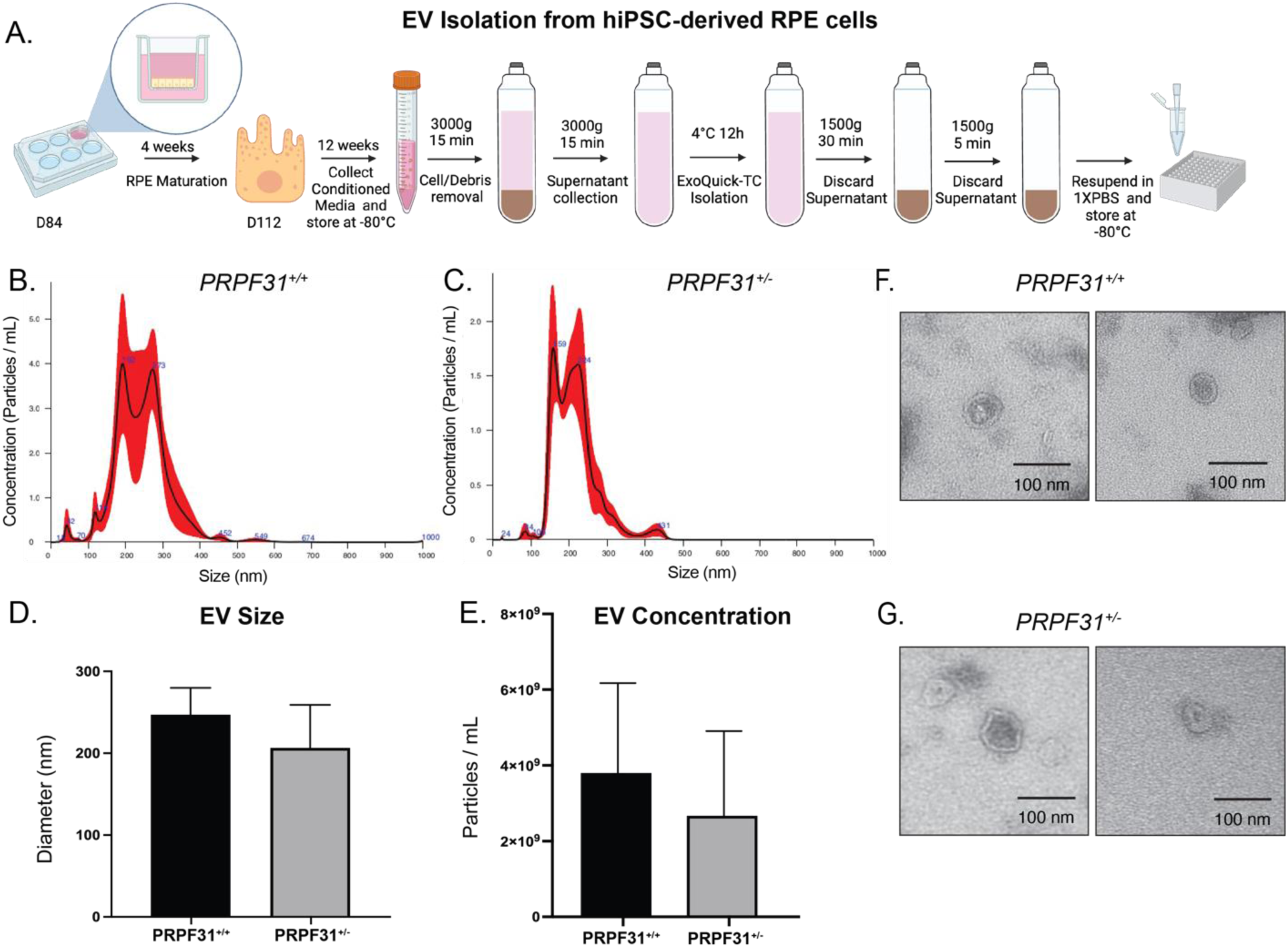
Characterization of EVs from PRPF31+/+ and PRPF31+/− iPSC-RPE cells. A-B) Distribution of EVs by size using nanoparticle tracking analysis technology. C) The average size of PRPF31^+/+^ EVs was 247 ± 16 nm and PRPF31^+/−^ EVs 206 ± 26 nm. D) The concentration of PRPF31^+/+^ EVs was 3.8x10^9^ ± 1.2x10^9^ EVs/ml and PRPF31^+/−^ EVs was 2.7x10^9^ ± 1.1x10^9^ EVs/ml. N=4, error bars: standard error of the mean. D) Ultrastructural analysis by TEM of EVs isolated from PRPF31^+/+^ and PRPF31^+/−^ conditioned media; scale bar is 100ϋm.

Characterization of the isolated EVs by nanoparticle tracking analysis (NTA) showed they ranged in size from 100nm to 400nm, and EVs from the *PRPF31*^+/-^ mutant and control RPE cells showed the same size distributions and concentrations (**Figure 2B-E**). Transmission electron microscopy (TEM) analysis showed the EVs were clear, rounded membrane vesicles as anticipated (**Figures 2 F-G**) ^61,62^.

### Sequencing of RNAs isolated from EVs produced by *PRPF31*-deficient and control RPE cells

To study the RNA contents of the EVs produced by the control *PRPF31*^+/+^ and mutant *PRPF31*^+/−^ hiPSC-RPE, we isolated EVs from the serum-free CM of 12 wells each of differentiated *PRPF31*^+/+^ and *PRPF31*^+/−^ iPSC-RPE cells grown in on Transwells as described above. Total RNA was isolated from the EVs and used to prepare both miRNA and poly-A RNA libraries from each of the 12 experimental and control cultures. The resulting set of 24 miRNA libraries was multiplexed to reduce batch effects and sequenced on an Illumina MiSeq instrument. At least 1.5 million reads were obtained for each miRNA library. Post QC, overall 8 mutants and 7 control samples were retained. For mutant samples total high quality (HQ) reads ranged from 1.5-2.6 million with a median of 1.9 million. For wild-type samples HQ reads ranged from 1.5-1.9 million with a median of 1.7 million.

Similarly, the set of 24 poly-A RNA libraries were multiplexed to reduced batch effects and sequenced on an Illumina NovaSeq instrument. Post QC, all 12 mutant and 12 wild-types samples were retained for differential gene expression analysis. For mutant samples HQ reads ranged from 46-120 million with a median of 67 million reads. For the wild-type samples, HQ reads ranged from 64-98 million with a median of 82 million reads.

### *PRPF31*^+/+^ and *PRPF31*^+/−^ EVs have distinct miRNA and poly-A RNA profiles

To investigate the effects of *PRPF31* haploinsufficiency and RPE dysfunction on the RNA contents of RPE EVs, we sequenced the miRNAs and poly-A-RNAs contained in EVs produced by control *PRPF31*^+/+^ and mutant *PRPF31*^+/−^ hiPSC-RPE as described above. Analyses of these data identified 135 miRNAs within the hiPSC-RPE EVs (**Supplementary Table S1**). To determine if any of these miRNAs could serve as a biomarker for RPE health, we compared the miRNA profiles in EVs from *PRPF31^+/+^* and *PRPF31^+/−^* hiPSC-RPE. Significant enrichment of miRNAs was determined using a combination of log fold change and FDR adjusted p-value (q-value). This differential enrichment analysis showed that 12 miRNAs were enriched in the EVs derived from the *PRPF31*^+/−^ hiPSC-RPE. In contrast, we found 6 miRNAs were depleted from the *PRPF31*^+/−^ hiPSC-RPE EVs (**Figure 3**). These miRNAs are also highlighted in **Supplementary Table S4**.

**Figure 3.**
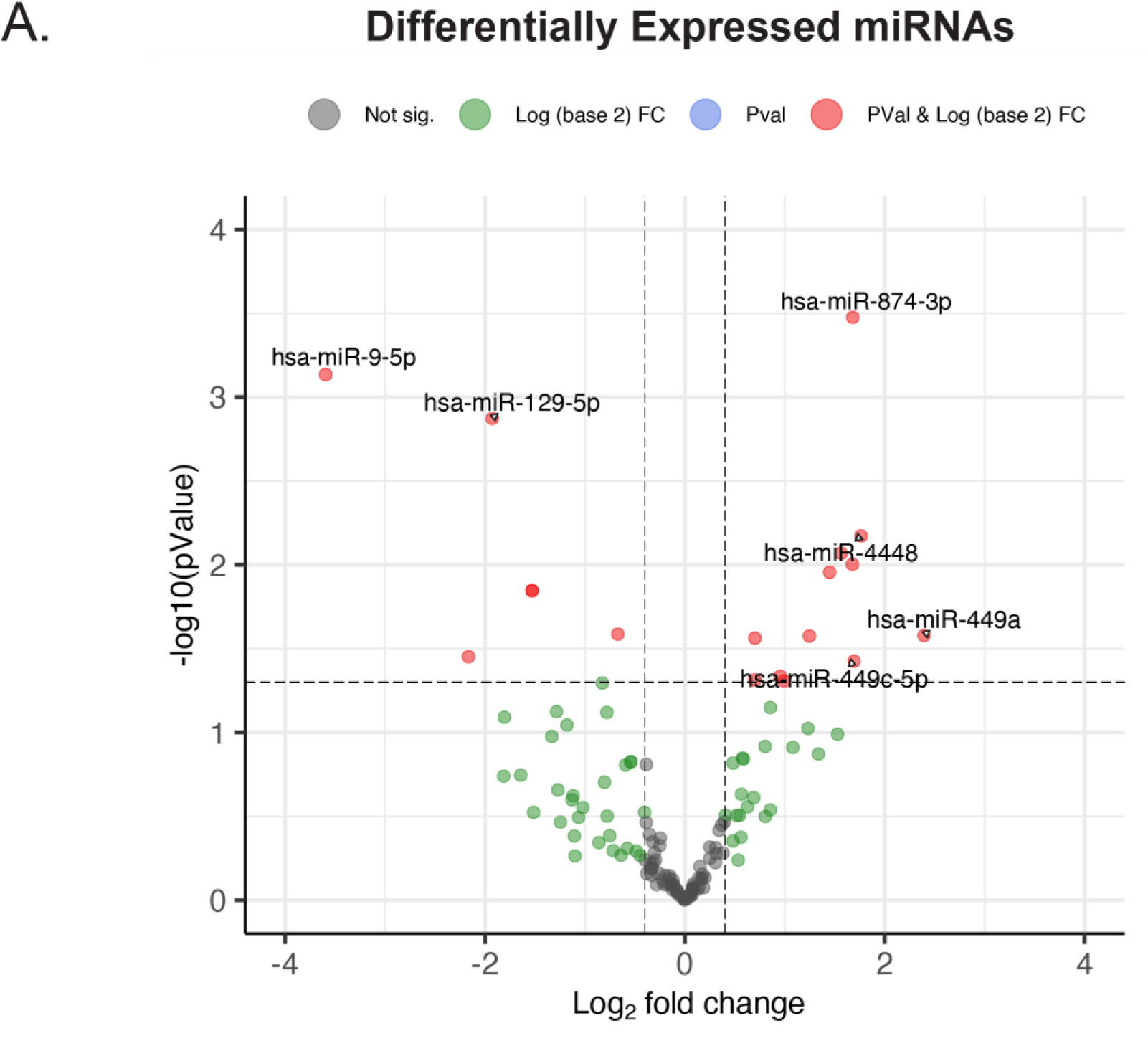
miRNAs profiling of EVs from PRPF31^+/+^ and PRPF31^+/−^. Volcano plots indicating the log2 (fold change) of differentially expressed miRNAs on the x-axis and the -log10 (adjusted P-value <0.05 8:7,Mut:WT) on the y-axis. miRNAs in red are significantly enriched (right) or depleted (left) in the PRPF31^+/−^ hiPSC-RPE EVs.

Examination of the poly-A RNA-seq reads mapped to the human genome revealed the presence of 31,776 poly-A RNAs in the hiPSC-RPE EVs (**Supplementary Table S2)**. Differential enrichment analysis of the poly-A RNA-seq data identified 865 differentially enriched poly-A RNAs, with 551 enriched in and 314 depleted from the *PRPF31^+/−^* hiPSC-RPE EVs (**Figure 4 and Supplemental Table S3**). Some of the poly-A RNAs were highly specific for *PRPF31*^+/−^ or *PRPF31^+/+^* hiPSC-RPE EVs, showing more than 1-million-fold enrichment in the respective EVs (log2 (1,000,000) = 19.9).

**Figure 4.**
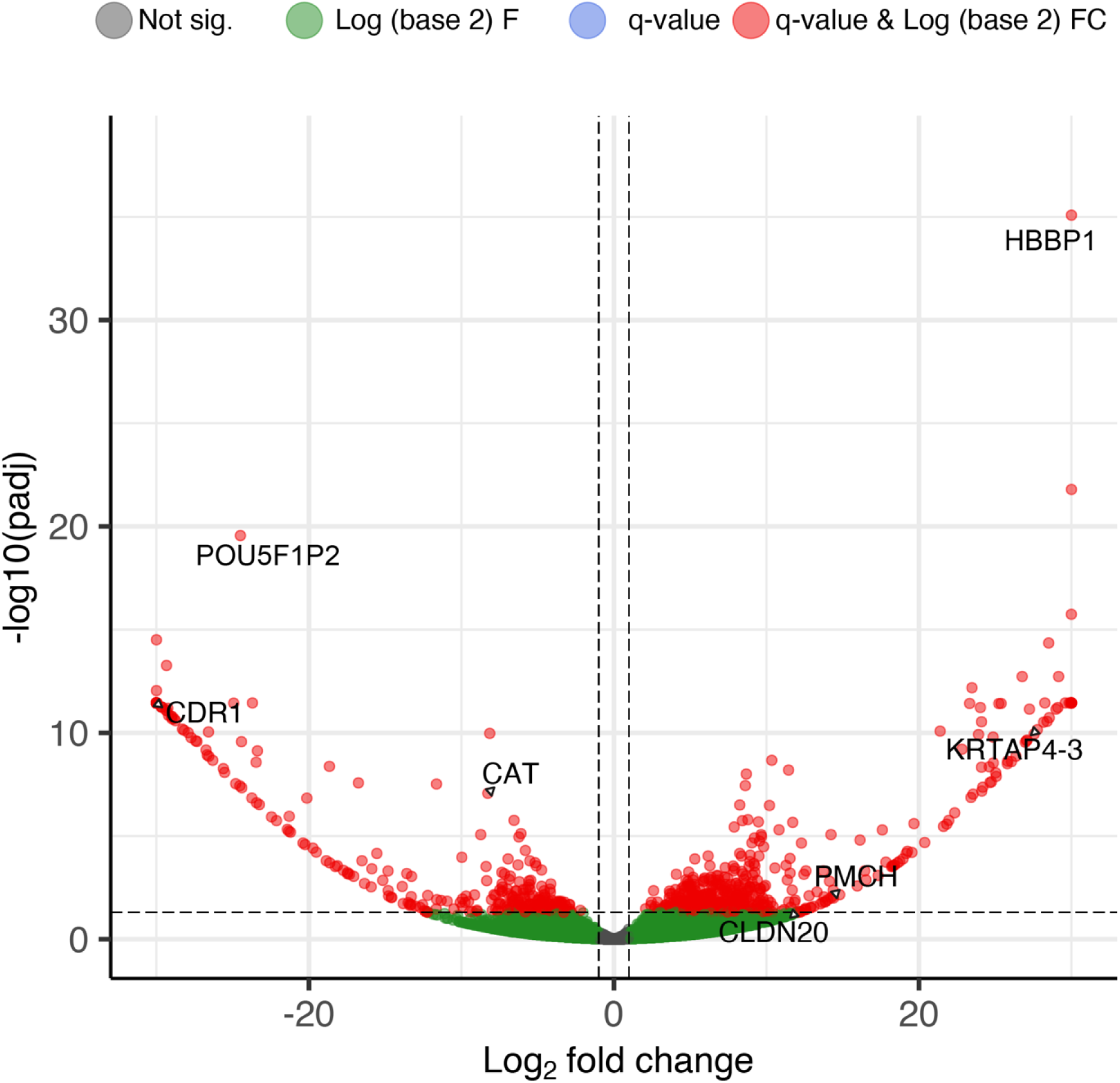
Comparison of the poly-A RNAs contained in EVs from PRPF31^+/+^ and PRPF31^+/−^ hiPSC-RPE. Volcano plot indicating the differentially expressed poly-A RNAs in the EVs isolated from PRPF31^+/−^ compared to the WT expressed as the log2 (fold change) on the x-axis and the -log10 (adjusted P-value <0.05, Mut:Wt;12:12) on the y-axis. Poly-A RNAs indicated by red dots were significantly enriched in (right) or depleted from (left) the PRPF31^+/−^ hiPSC-RPE based on fold change and adjusted P value.

### The majority of the miRNAs and poly-A RNAs contained in EVs are shared with hiPSC-RPE cells

To test the hypothesis that EV RNAs can be used as surrogate biomarkers for the health status of their source cells, we sequenced and compared the miRNAs and poly-A RNAs of the *PRPF31^+/+^*and *PRPF31^+/-^*hiPSC-RPE cells from which the EVs studied in the experiments described above were derived. These comparisons showed that the majority of miRNA and poly-A RNAs expressed in hiPSC-RPE cells are contained in the RPE-derived EVs. For miRNAs, 135 (64%) of the 211 miRNAs identified in the source hiPSC-RPE cells were present in the EVs isolated from the CM of those cells (**Supplementary Table S1 and S7**). EVs produced by the control *PRPF31^+/+^*hiPSC-RPE contained 90% (19,228) of the total 21,375 poly-A RNAs identified in the hiPSC-RPE (**Figure 5A**). Similarly, EVs produced by the mutant *PRPF31^+/−^* hiPSC-RPE contained 90% (19,247) of the 21,354 total poly-A RNAs identified in the mutant hiPSC-RPE (**Figure 5B and Supplementary Table S2 and S6**). These results indicate that the EVs produced by hiPSC-RPE cells contain the great majority of RNAs of their parent cells, supporting the potential use of EV RNAs as biomarkers. Interestingly, the EVs produced by the hiPSC-RPE cells also contained many additional poly-A RNAs not identified within their parent cells. *PRPF31^+/+^* EVs contained 11,984 unique poly-A RNAs and *PRPF31^+/−^* EVs contained 12,094 unique poly-A RNAs not found in their corresponding hiPSC-RPE cell transcriptomes (**Figure 5A-B**). The EVs also contained a greater percentage of lncRNAs and pseudogene mRNAs than the RPE cells (**Figure 5C-D**). Among the identified transcript RNA biotypes found in RPE cells, the predominant category consisted of protein-coding mRNAs (70%), followed by long non-coding RNAs (lncRNAs; 20%). In EVs the distribution of protein-coding mRNAs (54%), and lncRNAs (30%) analysis revealed a higher abundance of the latter. (**Figure 5C-D and Supplementary Table S2 and S6)**.

**Figure 5.**
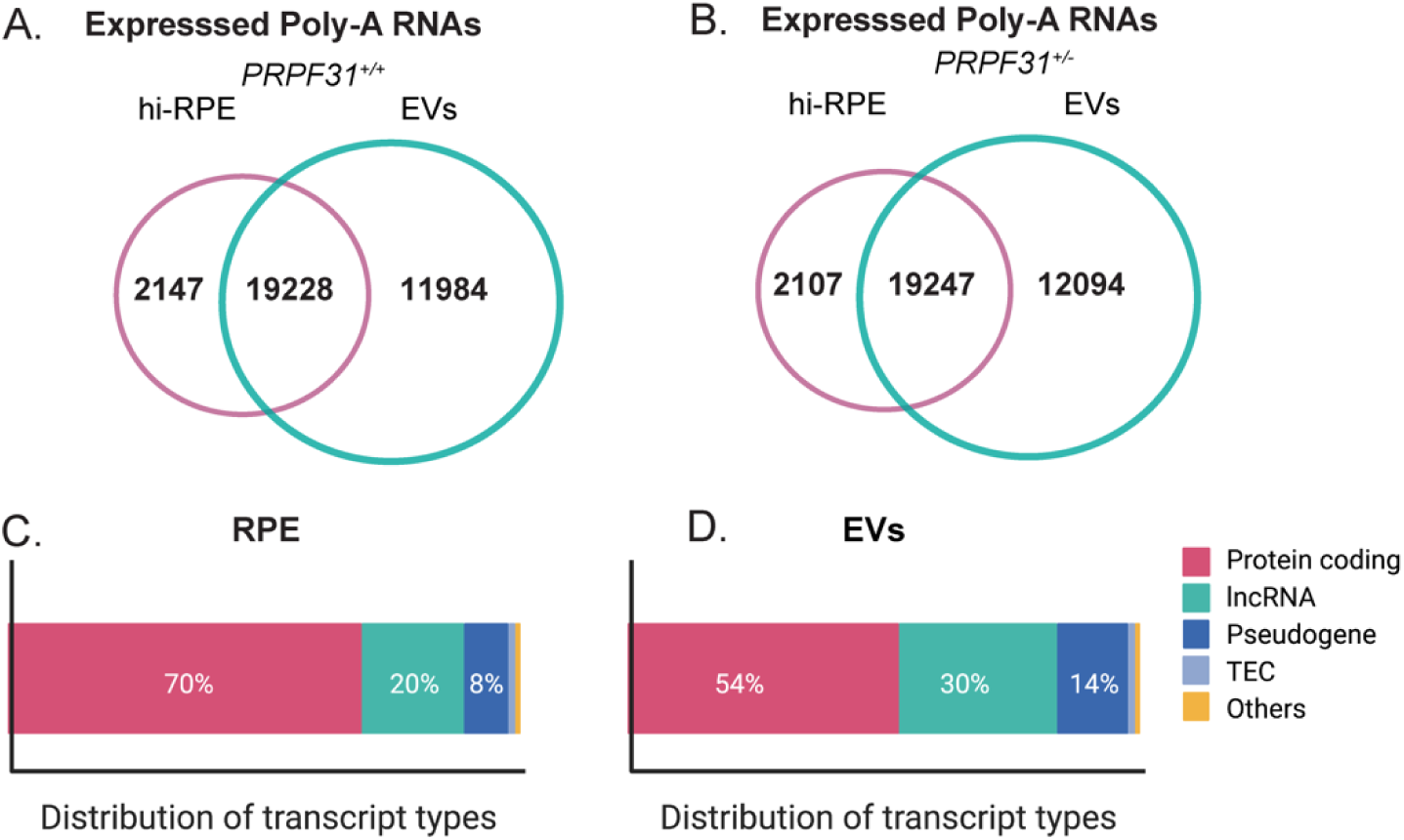
Comparison of RNAs identified in EVs produced by PRPF31^+/+^ and PRPF31^+/-^ hiPSC-RPE cells, and the RNAs contained in the hiPSC-RPE cells. (A, B) Venn diagrams showing unique and shared expressed poly-A RNAs among (A) PRPF31^+/+^ and (B) PRPF31^+/−^ hi-RPE cells and their EVs. (**C, D**) Relative biotype distribution of dysregulated transcripts in RPE cells and EV poly-A RNAs.

### Gene set enrichment analysis (GSEA) of poly-A RNAs and functional over-representation analysis (ORA) of miRNAs enriched and depleted in the *PRPF31* mutant

To explore the functional significance of the miRNAs and poly-A RNAs identified to be differentially enriched in the EVs derived from the *PRPF31*^+/-^ mutant hiPSC-RPE cells, we performed gene set enrichment analysis (GSEA) of poly-A RNAs and functional over-representation analysis (ORA) of miRNAs. GSEA for the 865 poly-A RNAs found to be differentially enriched or depleted in the EVs from *PRPF31*^+/-^ mutant hiPSC-RPE cells did not identify any enrichment of functional groups ^63^.

We used ORA to evaluate the potential functional significance of the miRNAs found to be differentially enriched in EVs from *PRPF31*^+/-^ vs. *PRPF31*^+/+^ hiPSC-RPE cells. ORA is used to evaluate the enrichment of transcripts regulated by miRNAs in functional categories derived from multiple databases, including gene ontology (GO, miRTarBase GO, miRWalk), target genes (miRTarBase), pathways (miRWalk, KEGG, miRPathDB), diseases (MNDR) and biological processes (miRPathD) ^55^. ORA found that 11 of the 18 miRNAs significantly enriched or depleted in the EVs derived from *PRPF31*^+/-^ mutant hiPSC-RPE cells were associated with eye diseases. Three of these miRNAs were associated specifically with retinal degeneration and 6 were associated with negative regulation of epithelial cell differentiation (**Supplementary Figure 4**).

Gene ontology analysis revealed that of the 18 miRNAs enriched or depleted in the EVs derived from *PRPF31*^+/-^ mutant hiPSC-RPE cells 6 were enriched in categories associated with cell de-differentiation or cell death and 6 with retinal function and degeneration (**Figure 6A and Supplementary Table S5**). For example, miRTarBase analysis identified *CDH1* and *RDH11* among the top most significantly overrepresented genes and each gene was targeted by a distinctive set of 6 miRNAs. *CDH1* is as a potential gene target of the differentially regulated miRNAs: miR-129-5p; miR-193b-3p; miR-34c-5p; miR-9-5p; miR-9-3p; miR-92a-3p. **(Figure 6B).** *CDH1* encodes for E-Cadherin, a structural protein found in the adherens junctions of the RPE cells which is dysregulated in pathological conditions of the RPE ^64,65^. Another distinctive set of differentially regulated miRNAs (miR-129-5p, miR-378a-3p, miR-34c-5p, miR-375-3p, miR-449b-5p, miR-449a) targeted *RDH11* gene which encodes Retinol dehydrogenase 11, related to visual phototransduction and with strong links with other RPE functions such as phagocytosis ^66–68^ **(Figure 6C).** The levels of *CDH1* and *RDH11* mRNAs were not altered in the *PRPF31*^+/-^ hiPSC-RPE cells themselves (**Supplementary Table S8**).

**Figure 6.**
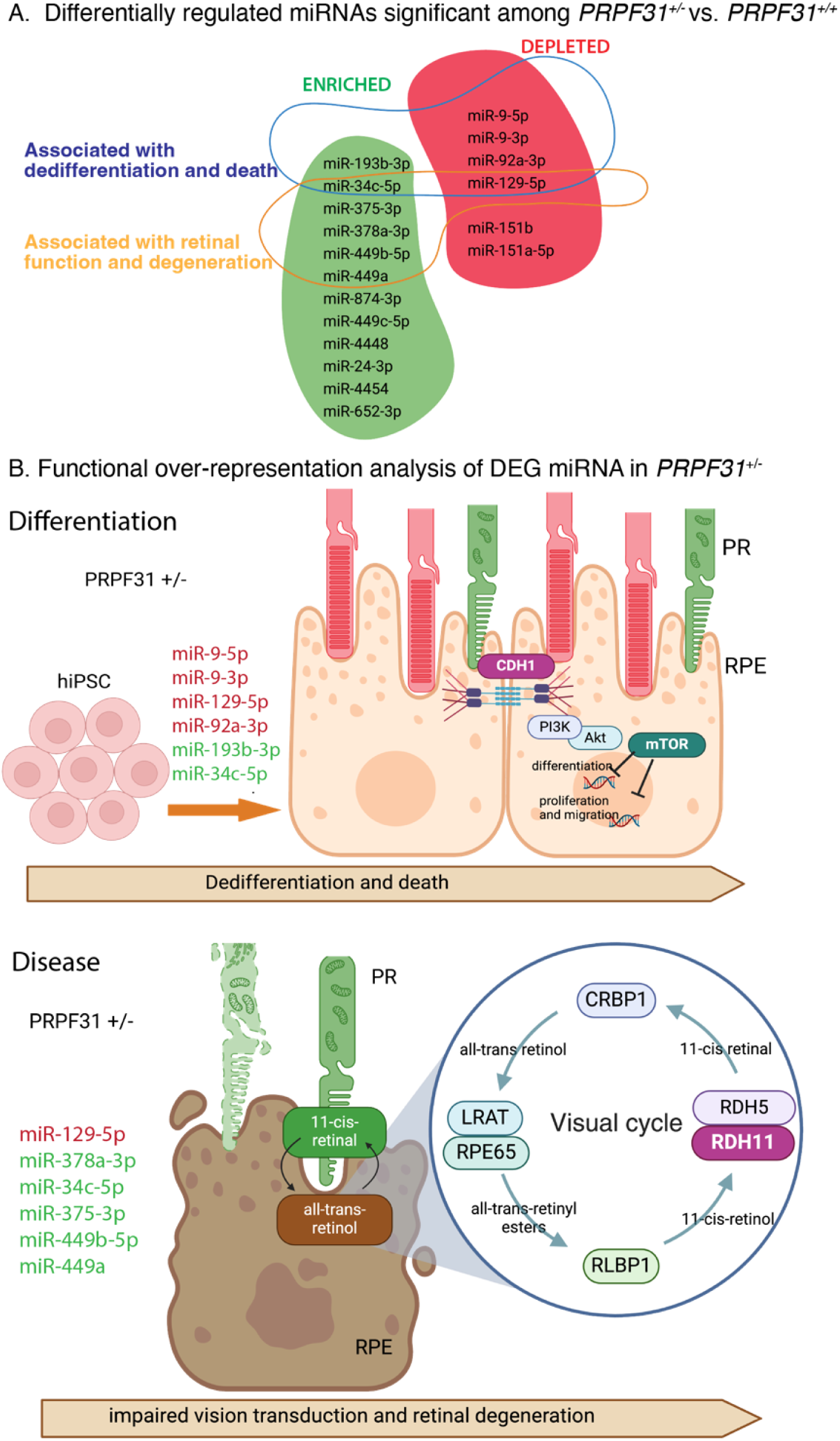
Functional over-representation analysis (ORA) of enriched and depleted miRNAs in the PRPF31 mutant. A) ORA analysis of differentially regulated miRNAs in PRFP31^+/-^ compared to PRFP31^+/+^ showed two distinctive subsets of miRNAs strongly associated with de-differentiation and cell death or associated with retinal function and degeneration. B. miRTarBase analysis identified CDH1 and RDH11 genes as top potential targets of the distinctive miRNAs subsets, which can explain the defects in the differentiation and function in the diseased PRPF31^+/-^ RPE cell.

### Validation of selected enriched poly-A RNAs and miRNAs by quantitative RT-PCR

We isolated total RNA from *PRPF31^+/+^* control and *PRPF31^+/−^* mutant hiPSC-RPE cells for qRT-PCR analyses of selected miRNAs and poly-A RNAs identified to be enriched or depleted in the EVs from the *PRPF31^+/−^* mutant hiPSC-RPE cells in the RNA sequencing studies. We choose 6 different poly-A RNAs among those differentially enriched in the *PRPF31^+/−^*hiPSC-RPE cells (**Supplementary Figure 1A**) compared to the *PRPF31^+/+^* control based on their read counts and RNA-seq trend. The qRT-PCR results showed that for 5 out 6 poly-A RNAs the levels in the EVs from *PRPF31^+/−^* and *PRPF31^+/+^* hiPSC-RPE cells were significantly different, and the changes detected were consistent with the RNA-seq data with regards to enrichment of depletion in the EVs from *PRPF31^+/−^* mutant hiPSC-RPE cells (**Supplementary Figure 2**). Similar qRT-PCR analyses were performed for 5 miRNAs identified to be differentially enriched in the EVs from *PRPF31^+/−^* mutant hiPSC-RPE cells from the RNA-sequencing data. qRT-PCR confirmed the differential enrichment of the miRNAs for 4 out of the 5 selected miRNAs in the *PRPF31*^+/−^ heterozygous and control EVs (**Supplementary Figure 3**).

## Discussion

We studied the RNA contents of EVs produced by control *PRPF31^+/+^*and mutant *PRPF31^+/−^* hiPSC-RPE cells to test the hypothesis that EV RNA contents have the potential to be used as surrogate biomarkers of RPE health. We have found that the RNA cargo in EVs mirrors that of their originating RPE cells, reflecting disease-induced changes in RPE cell differentiation and function. Further, by comparing the RNA profiles of EVs produced by *PRPF31^+/+^* and *PRPF31^+/−^* hiPSC-RPE cells, we found 18 miRNAs and 865 poly-A RNAs that are differentially enriched or depleted in the EVs from *PRPF31*^+/-^ mutant hiPSC-RPE cells. This includes many RNAs that were enriched or depleted at least 1-million-fold compared to EVs from control cells. Further, miRNAs differentially enriched in EVs from the *PRPF31*^+/-^ mutant hiPSC-RPE were identified as regulators of genes associated with RPE differentiation and function, supporting the functional relevance of the differentially enriched EV RNAs. Overall, these findings support the hypothesis that the RNA contents of EVs produced by RPE cells changes during disease, consistent with the potential use of EV RNAs as biomarkers of RPE health.

As a first step in assessing the potential use of EV RNAs as biomarkers of RPE health status, we compared the poly-A RNA profiles of EVs and their source cells. We found the majority (90%) of the poly-A RNAs detected in the parent hiPSC-RPE cells were contained in the EVs derived from those cells, supporting the concept that EV RNAs reflect the transcriptomes of the cells they are derived from ^29,35–39^. Of interest, we also found many poly-A RNAs were detected only in EVs, including additional lncRNAs. The higher abundance of lncRNA in the EVs is intriguing and suggests specific regulatory mechanisms at play. Long non-coding RNAs (lncRNAs) are thought to play regulatory roles in multiple biological processes, including the differentiation and function of RPE cells ^69,70^. Dysregulation of lncRNAs have been associated with age-related macular degeneration (AMD)^71,72^. Other studies have reported similar differences in RNA profiles and abundance when comparing EVs to their source cells ^73,74^. Although the mechanism of RNA sorting and loading into EVs is currently unknown, some studies suggest the contents of EVs are loaded by source cells as a mechanism of controlling intracellular availability of certain RNAs ^37,75,76^. Further studies will be needed to investigate the roles of EV lncRNAs in RPE biology and disease. It is also important to consider for this study and others, EVs were isolated from pooled CM that was collected for several weeks, while source cell RNA was isolated from one time point, so differences can also be due temporal RNA expression differences at the time of sample collection. According to the latest MISEV recommendations, no method for isolating EVs can completely guarantee the absence of non-vesicular particles. However, the primary potential contaminants, lipoprotein particles, do not contain RNA and, therefore, will not contribute to the RNA content in EVs for the purposes of this study.

Comparisons of the RNA contents of EVs produced by *PRPF31^+/+^*and *PRPF31^+/−^* hiPSC-RPE cells identified 12 miRNAs that were enriched and 6 miRNAs depleted in the EVs derived from the *PRPF31*^+/−^ hiPSC-RPE. Similarly, differential enrichment analysis of the poly-A RNA-seq data identified 551 poly-A RNAs enriched in and 314 depleted from the *PRPF31^+/−^* hiPSC-RPE EVs. Some of the poly-A RNAs were highly specific for *PRPF31*^+/−^ or *PRPF31^+/+^*hiPSC-RPE EVs, supporting their potential use as biomarkers of RPE health and/or disease. The RNAseq data comes from a large number of replicates (7-8 samples per condition) making these results statistically robust ^58^. Additionally, we have validated the directional change of RNA concentration for several miRNAs and poly-A RNAs using quantitative RT-PCR as an orthogonal method.

It has been previously demonstrated that mRNAs and miRNAs can be transferred via EVs and in turn regulate gene expression of recipient cells ^77–81^ The data presented here provide a starting point for future studies of the function(s) of RNAs contained in EVs produced by the RPE. For example, photoreceptor and RPE cells function together, with coordinated metabolism and recycling of shed photoreceptor outer segments via RPE phagocytosis ^82^. Since overall retinal function and health depends on these shared processes, it is attractive to hypothesize that EVs may be used for intercellular signaling between the RPE and photoreceptor cells, however further studies are needed to support this hypothesis ^83^.

We focused these initial studies of EV contents on RNAs since with current technologies better depth of sampling is possible for RNA sequencing than proteomic analysis ^84^. Further, EV RNAs have been reported to be promising biomarkers of cancer and other forms of neurodegeneration ^85–88^. Future studies could also evaluate the protein contents of RPE and retina derived EVs, as reported ^41^.

We were especially interested to find that some of the miRNAs differentially enriched in EVs from the *PRPF31*^+/-^ mutant hiPSC-RPE were identified as regulators of genes associated with RPE differentiation and function. Among the 18 most significantly altered miRNAs between the EVs produced by *PRPF31*^+/-^ and *PRPF31*^+/+^ cells, three were linked to retinal degeneration, and six were connected to the negative regulation of epithelial cell differentiation. This suggests that the defective differentiation and function of RPE cells caused by *PRPF31* haploinsufficiency is reflected in the miRNAs content in the EVs produced by the mutant RPE cells ^16,17^. We believe these findings support the hypothesis that EV RNAs can serve as surrogate biomarkers of retinal and RPE health status, and with further study may provide an alternative approach to assess the response of patients with IRDs to potential therapies, such as AAV-mediated gene therapies which show great promise for treating IRDs ^89^.

## Supporting information

Supplemental figures and descriptions

## Acknowledgements

We are grateful for the contributions of other Ocular Genomics Institute team members, including Daniela Pignatta and the Ocular Genomics Core for their assistance with the RNA-seq. We also acknowledge Philip Seifert and the Schepens Eye Research Institute Morphology Core for their assistance with the TEM imaging. We thank Amanda Fernández-Rodríguez for her assistance with miRNA sequencing analysis. The research reported was supported by a Stein Innovation Award from Research to Prevent Blindness (EAP), grants from the National Eye Institute to EAP (EY012910, EY020902), and a P30 Core Grant for Vision Research (EY014104).

## Abbreviations

AAV: adeno-associated virus
CM: conditioned media
EB: Embryoid Body
EVs: Extracellular vesicles
GSEA: gene set enrichment analysis
hiPSC: human induced pluripotent stem cell
hiPSC-RPE: hiPSC-derived RPE
IRD: inherited retinal degeneration
lncRNAs: long non-coding RNAs
MISEV: Minimal Information for Studies of Extracellular Vesicles
NTA: Nanoparticle tracking analysis
ORA: over-representation analysis
PRPF31: pre-mRNA processing factor 31
RPE: retinal pigment epithelium
RP: retinitis pigmentosa
TEM: transmission electron microscopy
ZO1: Zonula occludens

## Ethics approval and consent to participate

The work reported here has been acknowledged by the Mass General Brighma Instituttional Review Board as “not human subjects research” (approved application # 2022P000301)

## Availability of data and materials

All data supporting the findings of this study are available within the paper and its supplementary information. RNAseq raw sequence reads were deposisted in NCBIs SRA can be found in https://www.ncbi.nlm.nih.gov/bioproject/PRJNA824312.

## Authors’ contributions

All authors have made substantial contributions to this manuscript. EAP and MGH conception EAP HG and MGH design of the work; HG TK SM RFG EAP and MGH the acquisition, analysis, and interpretation of data; HG EAP and MGH have drafted the work and substantively revised it. All authors have approved the submitted version

