## Supplemental figures and descriptions for "The RNA content of extracellular vesicles from *PRPF31*^+/−^ hiPSC-RPE show potential as biomarkers of retinal degeneration"

### Supplementary Figures

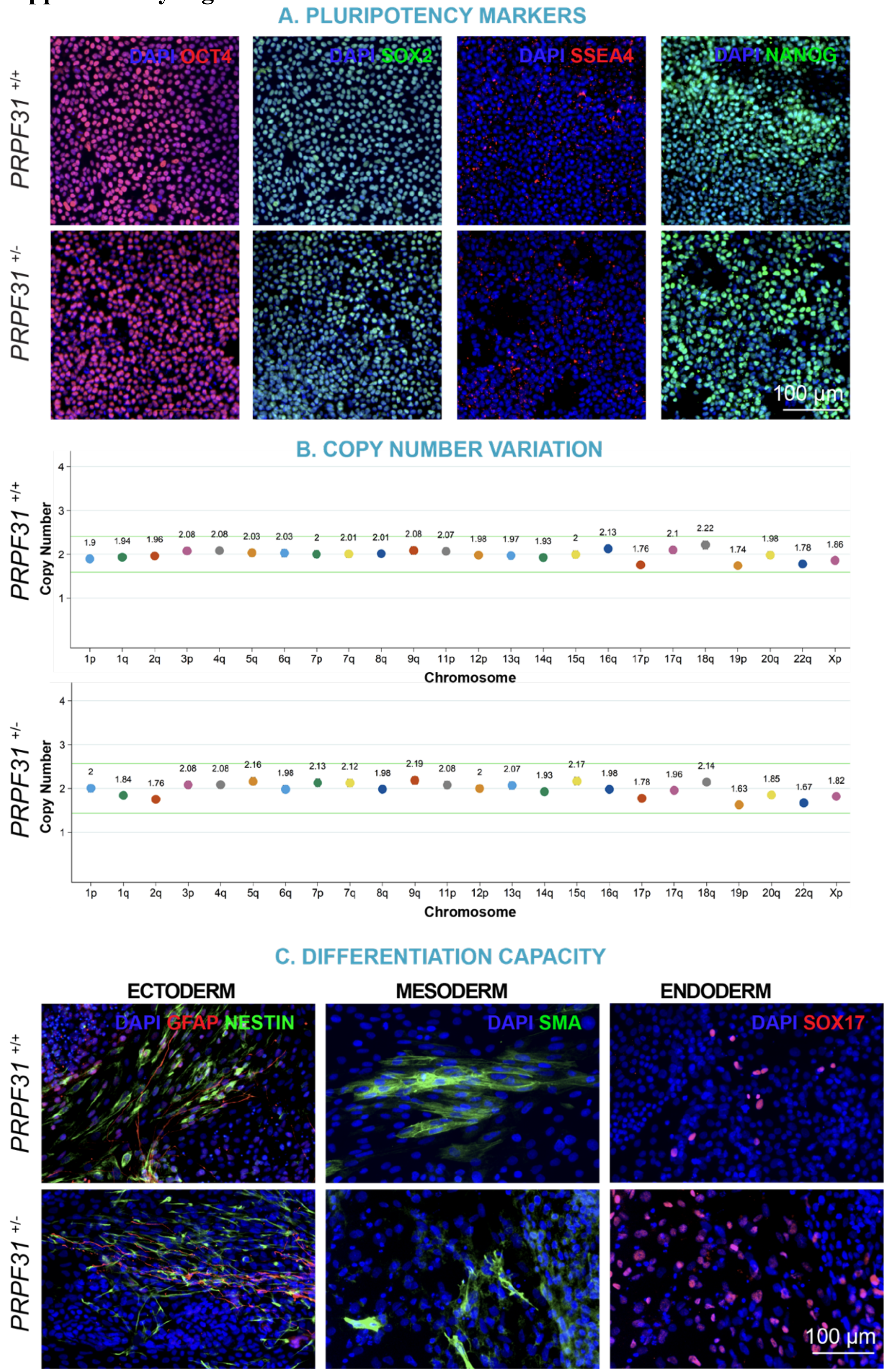

***Supplementary Figure 1. Characterization of CRISPR-Cas9 generated PRPF31$^{+/+}$ and PRPF31$^{+/-}$ hiPSC lines.*** A) Immunofluorescence analysis of hiPSC lines showing cells immunoreactive for pluripotency markers OCT4, SOX2, SSEA4 and NANOG. B) Copy number variation analysis showing absence of chromosomal abnormalities. C) Differentiation capacity of the *PRPF31*$^{+/+}$ and *PRPF31*$^{+/-}$ hiPSC lines using the embryoid bodies (EB) model. Immunofluorescence analysis of hiPSC-derived EBs depicting expression of the three germ layer markers: NESTIN, GFAP (ectoderm), SMA (mesoderm), and SOX17 (endoderm). Scale bar = 100 μm.

A. List of transcripts selectively expressed in EV isolated from RPE cells

| Gene ID | Gene Name | Gene Description | pValue | RNAseq trend |
|---|---|---|---|---|
| ENSG00000171217 | CLDN20 | claudin 20 [Source:HGNC Symbol;Acc:HGNC:2042] | 0.001305414 | UP |
| ENSG00000183395 | PMCH | pro-melanin concentrating hormone [Source:HGNC Symbol;Acc:HGNC:9109] | 9.76E-05 | UP |
| ENSG00000196156 | KRTAP4-3 | keratin associated protein 4-3 [Source:HGNC Symbol;Acc:HGNC:18908] | 1.68E-13 | UP |
| ENSG00000288642 | CDR1 | cerebellar degeneration related protein 1 [Source:HGNC Symbol;Acc:HGNC:1798] | 3.00E-15 | DOWN |
| ENSG00000240857 | RDH14 | retinol dehydrogenase 14 [Source:HGNC Symbol;Acc:HGNC:19979] | 1.96E-06 | DOWN |
| ENSG00000121691 | CAT | catalase [Source:HGNC Symbol;Acc:HGNC:1516] | 3.33E-10 | DOWN |

B. Fold change of selected poly-A RNA transcripts and p values obtained with qRT-PCR Analysis

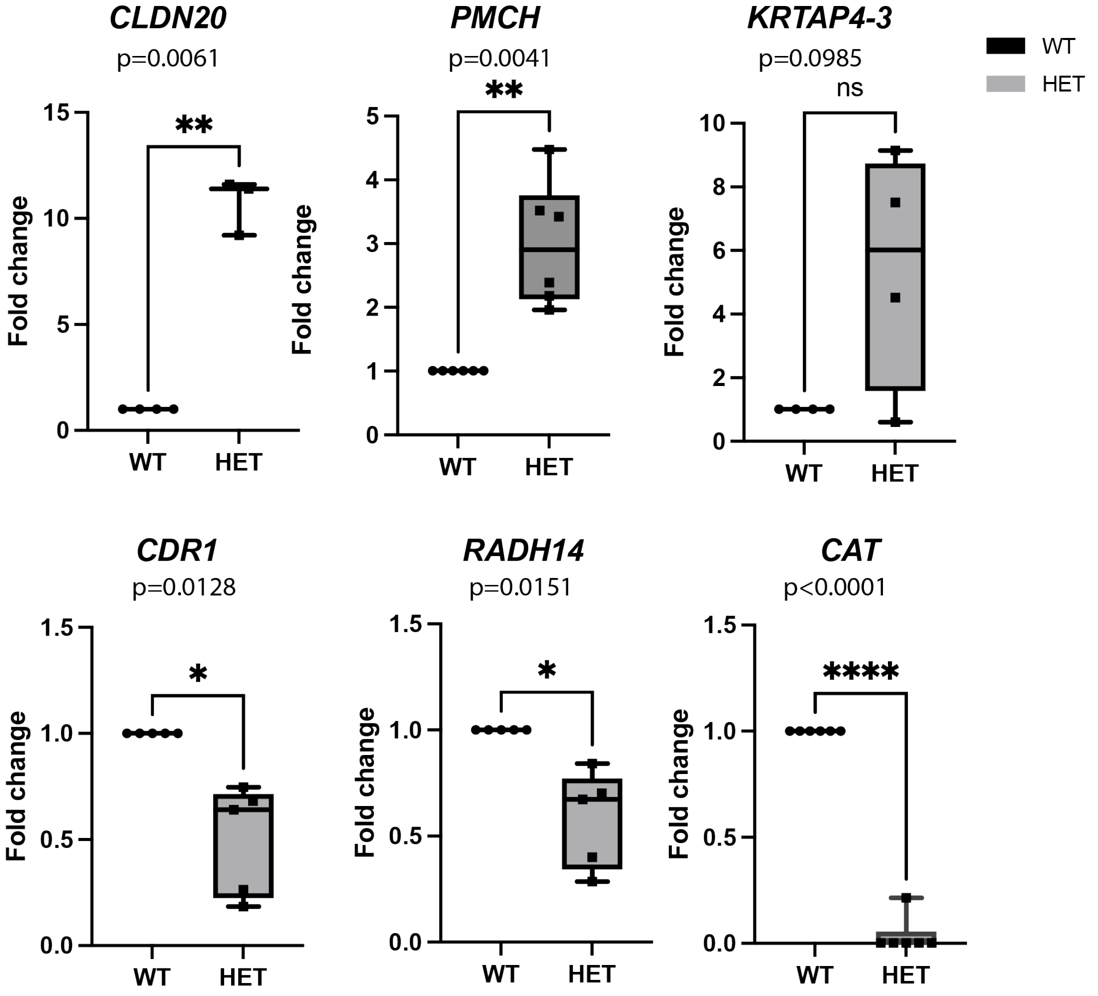

***Supplementary Figure 2. Validation of differentially regulated poly-A RNAs by qRT-PCR analysis.*** (A) List of poly-A transcripts selected to validate RNA sequencing expressed in either EVs from hiPSC-derived RPE from *PRPF31* wild type or mutant (B) Fold change values of the 5 most relevant genes expressed in hiPSC-derived RPE from *PRPF31* healthy or mutant. Fold change of *CLDN20, PMCH, KRTAP4-3, CDR1, RDH14* and *CAT* genes are presented as box-plots from the minimum to the maximum value, showing median (line) and mean. P-value of comparison of expression level of each gene within the two groups is reported. N=5 biological replicates were used for validation of RNA seq.

A. List of selected DEG  expressed in EV isolated from RPE cells

| ACCESSION miRBASE | miRNA Name | SEQUENCE | pValue | RNAseq trend |
|---|---|---|---|---|
| MIMAT0010251 | hsa-miR-449c-5p | UAGGCAGUGUAUUGCUAGCGGCUGU | 0.037543847 | UP |
| MI0016791 | hsa-miR-4448 | GGCUCCUUGGUCUAGGGGUA | 0.00671751 | UP |
| MI0001648 | hsa-miR-449a | UGGCAGUGUAUUGUUAGCUGGU | 0.026410276 | UP |
| MI0000490 | hsa-mir-206 | UGGAAUGUAAGGAAGUGUGUGG | 0.295261575 | DOWN |
| MI0000252 | hsa-miR-129-5p | CUUUUUGCGGUCUGGGCUUGC | 0.001340785 | DOWN |

B. Fold change of selected poly-A RNA transcripts and p values obtained with qRT-PCR Analysis

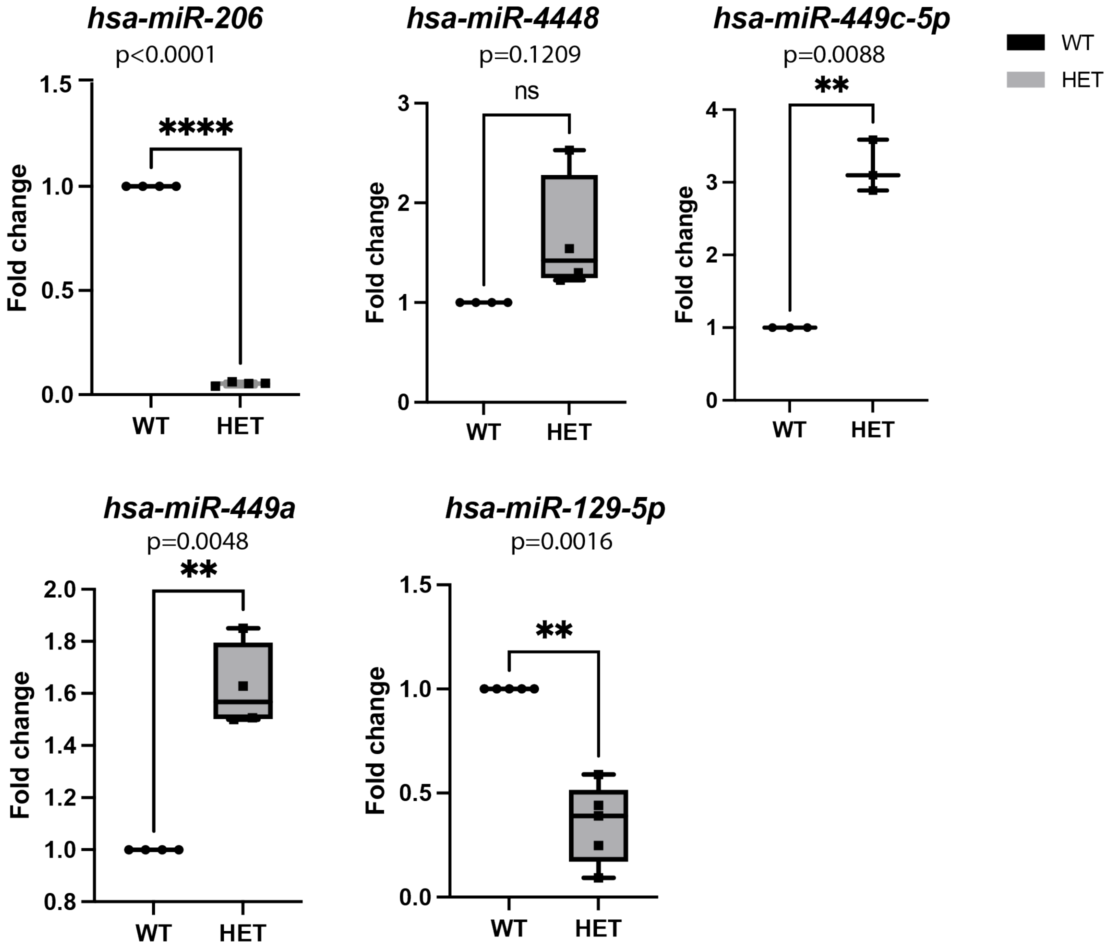

***Supplementary Figure 3. Validation of differentially regulated miRNA by qRT-PCR analysis.*** (A) List of selected miRNAs to validate RNA sequencing expressed in either EVs isolated from hiPSC-derived RPE from *PRPF31* wild type or mutant (B) Fold change values of the 5 most relevant miRNAs expressed in hiPSC-derived RPE from *PRPF31* healthy or mutant. Fold change of *hsa-miR-206*, *hsa-miR-4448, hsa-miR-449c-5p, hsa-miR-449a, hsa-miR-129-5p* miRNAs are presented as box-plots from the minimum to the maximum value, showing median (line) and mean. P-value of comparison of expression level of each gene within the two groups is reported. N=5 biological replicates were used for validation of RNA seq.

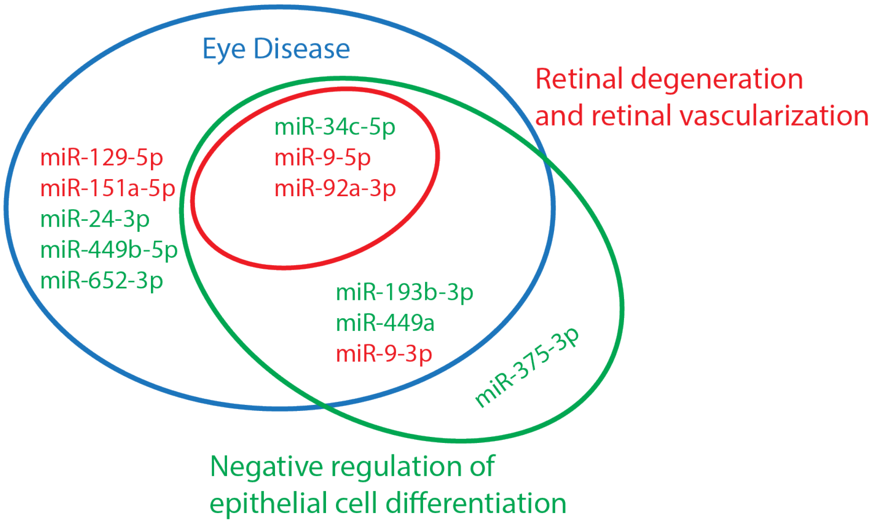

***Supplementary Figure 4. Functional over-representation analysis of DEG miRNAs in PRPF31$^{+/-}$*** depicting the miRNAs associated with the categories of Eye disease (blue), retinal degeneration and retinal vascularization (red) and negative regulation of epithelial cell differentiation (green). miRNAs showed in red font are depleted and in green font are enriched in the PRFP31 mutant.

**Supplementary Table Legends**

***Table S1. miRNAs identified in EVs***
Column A indicates the gene ID for the miRNAs detected.
Columns B and C show the read depth in average counts per million (CPM) of the miRNAs detected.

***Table S2. Poly-A RNAs identified in EVs***
Column A indicates the Ensembl gene ID for the poly-A RNAs identified in EVs
Column B indicates the gene name for the poly-A RNAs identified in EVs
Columns C and D indicates presence or absence of the gene in mutant or wildtype EVs respectively
Column E indicates the gene description
Column F indicates the gene biotype

***Table S3. Differentially enriched poly-A RNAs in EVs from mutant PRPF31 RPE cells***
Column A indicates the Ensembl gene ID for the poly-A RNAs differentially enriched in EVs from mutant RPE cells.
Column B indicates the log2FoldChange
Column C indicates the p-value
Column D indicates the p adjusted value
Column E indicates the gene name of the differentially enriched poly-A RNAs in EVs from mutant RPE cells.

Column F indicates the gene description of the differentially enriched poly-A RNAs in EVs from mutant RPE cells.
Column G indicates the gene biotype of the differentially enriched poly-A RNAs in EVs from mutant RPE cells.

***Table S4. Differentially enriched miRNAs in EVs from PRPF31 mutant RPE cells***
Column A indicates the number of rows
Column B indicates the miRNA id of the differentially regulated miRNA in EVs derived from mutant RPE cells. miRNAs depleted in the mutant are highlighted in bold and miRNAs enriched in the mutant are in regular font.
Column C indicates the miRNA accession number of the differentially regulated miRNAs in column A
Column D indicates the mature sequence of the differentially regulated miRNAs in column A
Column E indicates the precursor miRNA of the differentially regulated miRNAs in column A
Column F indicates the predicted gene targets of the differentially regulated miRNAs in column A according to the miRDB data base.

***Table S5. Over-representation analysis (ORA) of differentially enriched miRNAs identified from pairwise comparisons (PRF31 mutant vs. PRF31 wildtype)***
Column A indicates the miRNA database (e.g. target genes from miRTarbase), used to perform the ORA analysis
Column B indicates the subcategory (e.g. a certain target gene)
Column C indicates the type of enrichment
Column D indicates the p-value
Column E indicates the number of miRNAs that we would expect to find associated with each subcategory in column B.
Column F indicates the observed number of overrepresented miRNAs associated with each subcategory in column B according to the database indicated in column A.
Column G indicates the miRNAs or precursors from the differentially enriched test set that are contained in each subcategory in column B according to the database indicated in column A.

***Table S6. Poly-A RNAs identified in RPE cells***
Column A indicates the Ensembl gene ID for the poly-A RNAs identified in RPE cells
Column B indicates the gene name for the poly-A RNAs identified in RPE cells
Columns C and D indicates presence or absence of the gene in mutant or wildtype RPE cells respectively
Column E indicates the gene description
Column F indicates the gene biotype

***Table S7. miRNAs identified in hiPSC-RPE***
Column A indicates the gene ID for the poly-A RNA gene detected.
Columns B and C show the read depth in average counts per million (CPM) of the poly-A RNA gene detected.

***Table S8. Differentially enriched poly-A RNAs in mutant PRPF31 RPE cells***

Column A indicates the Ensembl gene ID for the poly-A RNAs differentially enriched in mutant RPE cells.
Column B indicates the log2FoldChange
Column C indicates the p-value
Column D indicates the p adjusted value
Column E indicates the gene name of the differentially enriched poly-A RNAs in mutant RPE cells.
Column F indicates the gene description of the differentially enriched poly-A RNAs in mutant RPE cells.
Column G indicates the gene biotype of the differentially enriched poly-A RNAs in mutant RPE cells.

***Table S9. TaqMan Gene Expression Assays for validation of poly-A RNA expression by qRT-PCR***
Column A indicates the Ensembl gene ID for the poly-A RNAs selected for qRT-PCR analysis
Column B indicates the gene name for the poly-A RNAs selected for qRT-PCR analysis
Column C indicates the gene description
Column D indicates the Taqman Gene Expression Assay ID

***Table S10. TaqMan Advanced miRNA Assay for validation of miRNA expression by qRT-PCR***
Column A indicates the miRNA Accession for the miRNAs selected for qRT-PCR analysis
Column B indicates the miRNA ID
Column C indicates the Taqman miRNA Assay ID
